# Craniofacial heterochrony in marsupials involves dynamic changes to cellular and molecular mechanisms underlying neural crest migration

**DOI:** 10.1101/2025.09.23.678178

**Authors:** Axel H Newton, Ella R Farley, Andrew T Major, Jennifer C Hutchison, Ben M Lawrence, Yoshio Wakamatsu, Karen E Sears, Marilyn B Renfree, Aidan M C Couzens, Geoff Shaw, Sara Ord, Richard A Schneider, Andrew J Pask

## Abstract

The neural crest is a vertebrate innovation central to craniofacial development and evolution. While the gene regulatory networks guiding neural crest development are well characterized, the mechanisms generating species-specific craniofacial diversity remain poorly understood. Marsupials provide a unique model for studying neural crest plasticity, having evolved accelerated patterns of craniofacial development. This adaptation arises because marsupials are born altricial after a short gestation yet require well-developed mouthparts to attach to a teat and continue development in the pouch. However, how marsupials achieve this heterochronic shift in neural crest development is largely unknown. Here we investigate the cellular and molecular mechanisms underlying their distinct heterochrony across three marsupial species, with mouse and chicken as comparators. We reveal that marsupials produce dense pre-migratory aggregates of neural crest cells at the borders of a still-flat, pre-neurulation embryo, which emigrate as cohesive, E-cadherin-positive, vimentin-negative epithelial-like sheets, potentially facilitating rapid establishment of the facial prominences. Re-analysis of published marsupial single-cell data further suggests that the mesenchymal transcriptional programme is deployed after delamination and migration. These cellular behaviours have not been described in other amniotes, instead resembling patterns in anamniotes which similarly exhibit accelerated craniofacial development to support early feeding. Marsupials thus appear to have evolved a similar mechanism of neural crest migration to facilitate their developmental heterochrony. These findings suggest that vertebrate neural crest migration may be shaped by life history as well as phylogeny, providing new perspectives on neural crest plasticity and the developmental mechanisms driving craniofacial diversity.

**Significance statement:** Newborn marsupials resemble embryos, yet must crawl unaided to the teat and suckle within minutes of birth. This demands a functional face far earlier than in other mammals. We find that the cells that build that face, the cranial neural crest, behave unlike those of any other amniote examined: they gather into dense clusters at the edges of a still-flat embryo and leave as cohesive sheets that retain their attachments to one another, rather than dispersing as single cells as they do in mice and chickens. Frogs and fishes, which also feed at an immature stage, do much the same. How neural crest cells migrate therefore appears to track the demands of life history, not ancestry alone.

## Introduction

Cranial neural crest cells are migratory, multipotent progenitors which generate the bone, cartilage, connective tissue and musculature of the vertebrate face *(1)*. Their migration from the dorsal neural tube to populate the facial prominences is one of the most studied morphogenetic events in vertebrate development *(2, 3)*, and the gene regulatory network (GRN) governing their induction, specification and delamination is deeply conserved (*4–11*). However, despite this conservation, the cellular mechanics of neural crest migration vary substantially across taxa *(12, 13)*. For example, in studied amniotes such as birds, reptiles and eutherian mammals, cranial neural crest cells delaminate during neural tube closure (neurulation), undergoing an epithelial-to-mesenchymal transition (EMT), remodelling their cadherin repertoire, acquiring vimentin, and dispersing as mesenchymal cells through the extracellular matrix (*14–21*). In contrast, cranial neural crest cells in anamniotes such as amphibia and ray-finned fish emigrate from the flattened neural plate epithelium as dense, epithelial-like sheets, collectively migrating and retaining cell-cell contacts and cohesive group behaviour during the initial phase of migration (*22–31*). While these distinct patterns generally follow the amniote–anamniote boundary, a major exception can be seen within marsupial mammals. Here, cranial neural crest cells emigrate before neural fold apposition and neural tube closure, from a still-flattened neural plate epithelium, more closely resembling the anamniote condition (*32–31*). This variation raises the question of whether the evolution of neural crest migratory mode is fixed by phylogeny or instead shaped by other constraints.

Marsupial mammals are well known for their unique mode of development with altricial birth and prolonged postnatal growth. Born in an otherwise embryonic state after a short gestation, altricial neonates must then immediately crawl to the teat and begin suckling. This places acute functional demands on orofacial structures, leading to markedly precocial craniofacial development (*37–44*). This extends to the underlying cranial neural crest cells, where marsupials have evolved a heterochronic shift in when cranial neural crest cells form, emigrate and populate the facial prominences, both relative to overall embryonic stage and other studied amniotes (*32–34*). Pioneering observations by comparative embryologists J. P. Hill and K. P. Watson noted that this early emigration appeared to occur from a flat, pre-neurulation embryo, leading Hill to remark in a letter: *“ the flat somiteless embryo*…*is the most interesting not to say puzzling and in some things, I think unique. I verily begin to believe the head mesoderm is derived from the ectoderm!” (32)*. Modern molecular work has confirmed these early observations, identifying evolutionary changes to expression and regulation of SOX9 as a contributor to precocious neural crest specification in the opossum *Monodelphis* domestica *(45, 46)*. Additionally, a single-cell transcriptomic atlas of *Monodelphis* organogenesis confirmed that anterior structures, including the neural crest, initiate and progress through their transcriptional programs earlier and faster than in eutherians *(47)*. Yet whether marsupial cranial neural crest heterochrony is accompanied by morphogenetic changes to formation and migration remains largely unaddressed.

In this study we show that marsupial cranial neural crest cells depart from canonical amniote morphogenetic processes, instead possessing similarities to those observed in anamniotes. Marsupial neural crest cells first form dense pre-migratory aggregates at the neural plate borders of the flattened neural plate epithelium, before delaminating and migrating as E-cadherin-positive epithelial-like sheets, displaying characteristics consistent with collective migration, to rapidly sculpt and fill the facial primordia. Re-analysis of *Monodelphis* embryo single-cell data *(47)*, suggests that the transcriptional EMT programme is deployed after delamination, rather than before it, inverting the canonical amniote sequence. Marsupial cranial neural crest development thus appears to represent an exception to the canonical amniote strategy, displaying migratory characteristics convergent with anamniotes. Furthermore, we show that the tempo of cranial neural crest formation and protrusion of the facial prominences is consistent with the altriciality of marsupial species at birth, linking neural crest development to functional feeding constraints. These data suggest that neural crest migration mode may be shaped by life history constraints rather than phylogenetic relatedness alone. The functional demands of early feeding may have independently favoured similar neural crest migration strategies across vertebrate lineages as divergent as marsupials and frogs.

## Results

### Marsupial cranial neural crest cells form dense pre-migratory aggregates within the neural plate borders

Owing to the scarcity of marsupial embryos, all panels are representative single experiments without replicates. To characterise the morphogenetic basis of heterochronic craniofacial development in marsupials, we first examined cranial neural crest cell formation during neurulation, a stage when the planar embryo displays visibly raised and protruding ‘headfolds’ *(48, 49)*. Wholemount fluorescent imaging of the neural crest specification and migration markers SOX9, SNAI2, TFAP2A and SOX10 in *Sminthopsis crassicaudata* embryos revealed that the ‘headfolds’ were the TFAP2A-positive neural plate borders, densely packed with SOX9, SNAI2, SOX10-positive cranial neural crest cells (Figure 1A,B). These protruded dorsally as thickened aggregates from the otherwise flat embryo (Figure 1B,B’), marking the future frontonasal process (FNP) and first pharyngeal arch (PA1), with similar patterns to *Monodelphis (45)*.

**Figure 1.**
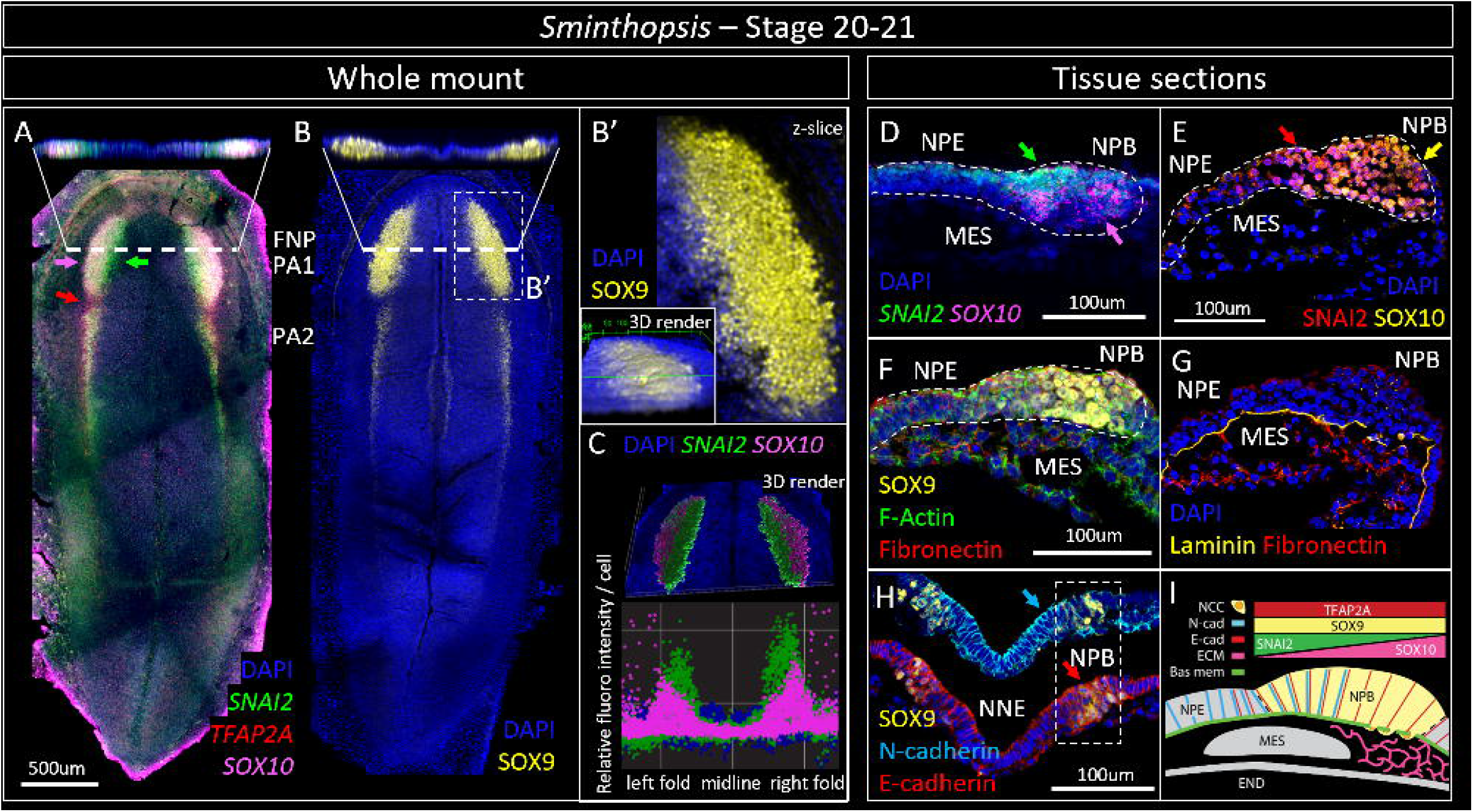
Marsupial cranial neural crest cells form dense pre-migratory aggregates at the neural plate border. Cranial neural crest formation in *Sminthopsis* embryos at McCrady stage 20–21. **A)** Wholemount *SNAI2, TFAP2A* and *SOX10* expression and **B⍰)** SOX9 protein localisation, showing dense headfolds of presumptive pre-migratory neural crest cells within the neural plate borders demarcating the presumptive frontonasal process (FNP) and first pharyngeal arch (PA1); transverse optical z-slices (top strips) show the headfolds protruding dorsally from the otherwise flat embryo. Bill) Magnified z-slice and 3D render of the SOX9-positive headfold. **C)** 3D volumetric render and per-cell quantification of A) showing relative SNAI2 and SOX10 fluorescence intensity, revealing a mediolateral SNAI2:SOX10 gradient across the neural plate borders (midline to left and right folds). **D–H)** Transverse tissue sections through the cranial neural crest domain: **D**,**E)** SNAI2 and SOX10 mark a densely packed, multilayered aggregate at the neural plate border (NPB), distinct from the neural plate epithelium (NPE) and underlying mesoderm (MES); **F)** SOX9, F-actin and fibronectin; **G)** laminin and fibronectin show the basement membrane remains intact beneath the aggregate; **H)** SOX9-positive aggregate cells retain E-cadherin-positive junctions but lack N-cadherin at their leading edges (NNE = non-neural ectoderm). **I)** Summary schematic of the pre-migratory aggregate, showing the relative distributions of TFAP2A, SOX9, SNAI2, SOX10, N-cadherin and E-cadherin over an intact basement membrane prior to delamination. NPE, neural plate epithelium; NPB, neural plate border; MES, mesoderm; END, endoderm.

Tissue sections through the cranial neural crest domains confirmed the aggregates were comprised of densely packed, multilayered cells positive for SOX9, SNAI2 and SOX10 (Figure 1D–F). Interestingly, these cells appeared to be uniformly SOX9 positive (Figure 1B,F), as in *Monodelphis (45)*, but displayed a mediolateral gradient of SNAI2 and SOX10 expression (Figure 1A,D). To quantify this internal organisation of cells within the aggregates, we examined the relative fluorescence intensity of *SNAI2* and *SOX10* per cell across the mediolateral axis of the aggregates, using 3D volumetric rendering of confocal stacks (Figure 1C). This confirmed that the aggregates were internally organised with a mediolateral gradient of *SNAI2:SOX10* expression, with SNAI2-high/SOX10-low cells in the medial portion of the neural plate border, but SNAI2-low/SOX10-high cells at the lateral leading edges of the neural plate borders. A comparable lateral-high/medial-low gradient of PAX7 has been reported within the pre-migratory cranial neural crest domain of *Monodelphis (45)*. This gradient suggests that SOX10-positive cells at the lateral margins of the aggregates are primed for delamination and breakdown of the basement membrane.

Critically, these cell aggregates had not yet initiated migration into the extracellular matrix, as laminin staining confirmed the basement membrane remained intact beneath (Figure 1G), and immunostaining for E-cadherin and N-cadherin revealed that the SOX9-positive cells retained a coherent epithelial morphology with strong E-cadherin-positive junctions and comparatively weak N-cadherin at their lateral edges (Figure 1H). Together, these data revealed that marsupial cranial neural crest cells initially form as dense pre-migratory aggregates within the neural plate borders, consistent with a model where aggregates are not uniformly pre-migratory but already establishing the cellular hierarchy that will drive subsequent migration. Given the requirement for rapid patterning and formation of the marsupial facial prominences (32–34), the cranial neural crest aggregates may serve as a reservoir of pre-migratory cells (Figure 1I).

### Marsupial cranial neural crest cells show early activation of the gene regulatory network

To identify molecular mechanisms underlying this heterochronic aggregate formation, we examined neural crest gene markers at the primitive streak stage — when *SOX9, BMP2* and *PAX7* can first be detected in *Monodelphis (45)*. Wholemount fluorescent imaging in primitive streak stage 19 *(49, 50) Sminthopsis* and *Monodelphis* embryos compared the pluripotency gene POU5F1 against the neural crest specifiers *FOXD3, SOX9* and TFAP2A, and delamination and migration markers *SNAI2* and *SOX10* (Figure 2A–D). While *POU5F1* predictably labelled the epiblast of both species, *SOX9* and *FOXD3* were additionally co-expressed within the forming neural plate epithelium, and *TFAP2A* was present in the neural plate borders (Figure 2A-E), suggesting priming of neural crest specification. Indeed, paired domains of *SOX10* were identified at the neural plate borders of stage 20 *Monodelphis* embryos (Figure 2E,E’) – a marker of pre- and post-migratory neural crest cells *(28)*. Early co-activation of core neural crest specifier genes within primitive streak stage embryos suggests the marsupial neural plate borders are primed for heterochronic cranial neural crest formation. This compression of the cranial neural crest GRN timeline is consistent with, but does not by itself demonstrate, a molecular mechanism underlying the precocious accumulation of cranial neural crest cells *(45, 46)*.

**Figure 2.**
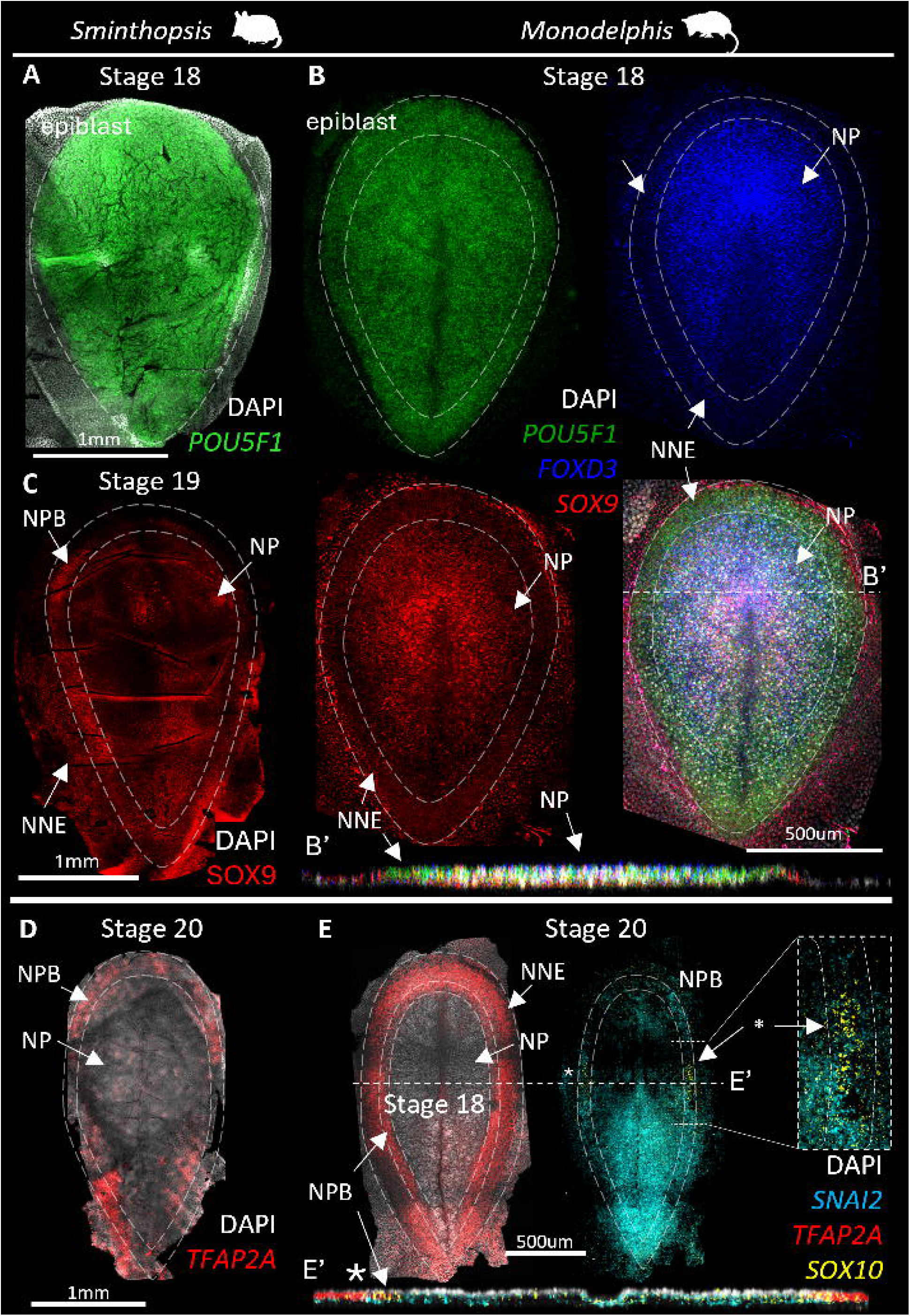
Neural crest specification within the neural plate borders of marsupial embryos. Wholemount visualisation of pluripotency, neural plate border and neural crest specification genes. **A)** *POU5F1* marks the epiblast of the *Sminthopsis* primitive-streak (stage 18) embryo. **B)** In the *Monodelphis* stage 18 embryo, *POU5F1, FOXD3* and *SOX9* delineate the boundary between the neural plate (NP) and non-neural ectoderm (NNE), with *FOXD3* and *SOX9* restricted to the forming neural plate in wholemount and optical z-section **(B⍰). C**,**D)** In the elongating stage 19-20 *Sminthopsis* embryo, SOX9 becomes restricted to the neural plate borders (NPB) alongside TFAP2A. **E)** In the stage 20 *Monodelphis* embryo, *SNAI2* is restricted to the underlying mesoderm *(45)*, while unexpectedly, *SOX10* showed paired expression domains at the neural plate borders (asterisks; magnified inset and z-section, **E⍰)**, suggesting that pre-migratory neural crest cells are first specified in the flattened, late primitive-streak-stage marsupial embryo. NP, neural plate; NNE, non-neural ectoderm; NPB, neural plate border.

### Marsupial cranial neural crest cells emigrate as epithelial-like sheets

With precocious cranial neural crest specification and formation in marsupials established, we next looked to determine the fate and migratory characteristics of the cell aggregates. Wholemount SOX9, SNAI2 and SOX10 immunostaining across stage 22–24 *Sminthopsis* embryos *(48)* revealed that marsupial cranial neural crest cells rapidly migrate as a densely packed wave to form the facial prominences (Figure 3A-C), with separation of the FNP, maxillary and mandibular prominences by stage 24. Similar dense sheets of SOX10+ NCCs are observed in the forming facial prominences of *Monodelphis* embryos (Figure S1) *(45)*. Tissue sections through the developing head revealed that migratory cells maintained the SNAI2:SOX10 mediolateral gradient, with newly migrating cells repressing SNAI2 and maintaining SOX10 in leader cells at the migratory front (Figure 3A’,B’). Furthermore, the densely packed migratory neural crest cells appeared to move as a single sheet through minimal ECM, wedged as a coherent mass between the surface ectoderm and the underlying endoderm and mesoderm (Figure 3A’-C’). Additional immunostaining for laminin and fibronectin revealed that leading cells appeared to navigate above the fibronectin extracellular matrix along the laminin basement membrane of the surface ectoderm (Figure 3D). In addition, staining for cell-cell junction and cytoskeletal proteins E-cadherin, N-cadherin and vimentin revealed that the densely packed migratory cells retained strong E-cadherin cell-cell junctions, lacked N-cadherin, and showed no detectable vimentin (Figure 3A“,B”). Importantly, both markers labelled adjacent tissues within the same sections and under identical acquisition settings: vimentin was detected in mesenchymal cells immediately beneath and lateral to the SOX10-positive domain, and N-cadherin strongly labelled the adjacent neuroepithelium and surface ectoderm (Figure 3A⍰⍰,B⍰⍰, arrows). The absence of signal within the neural crest population therefore reflects a genuine difference in protein localisation rather than a failure of antibody cross-reactivity in marsupial tissue. These morphological characteristics are consistent with collective sheet migration and a partial EMT phenotype (Figure 3E).

**Figure 3.**
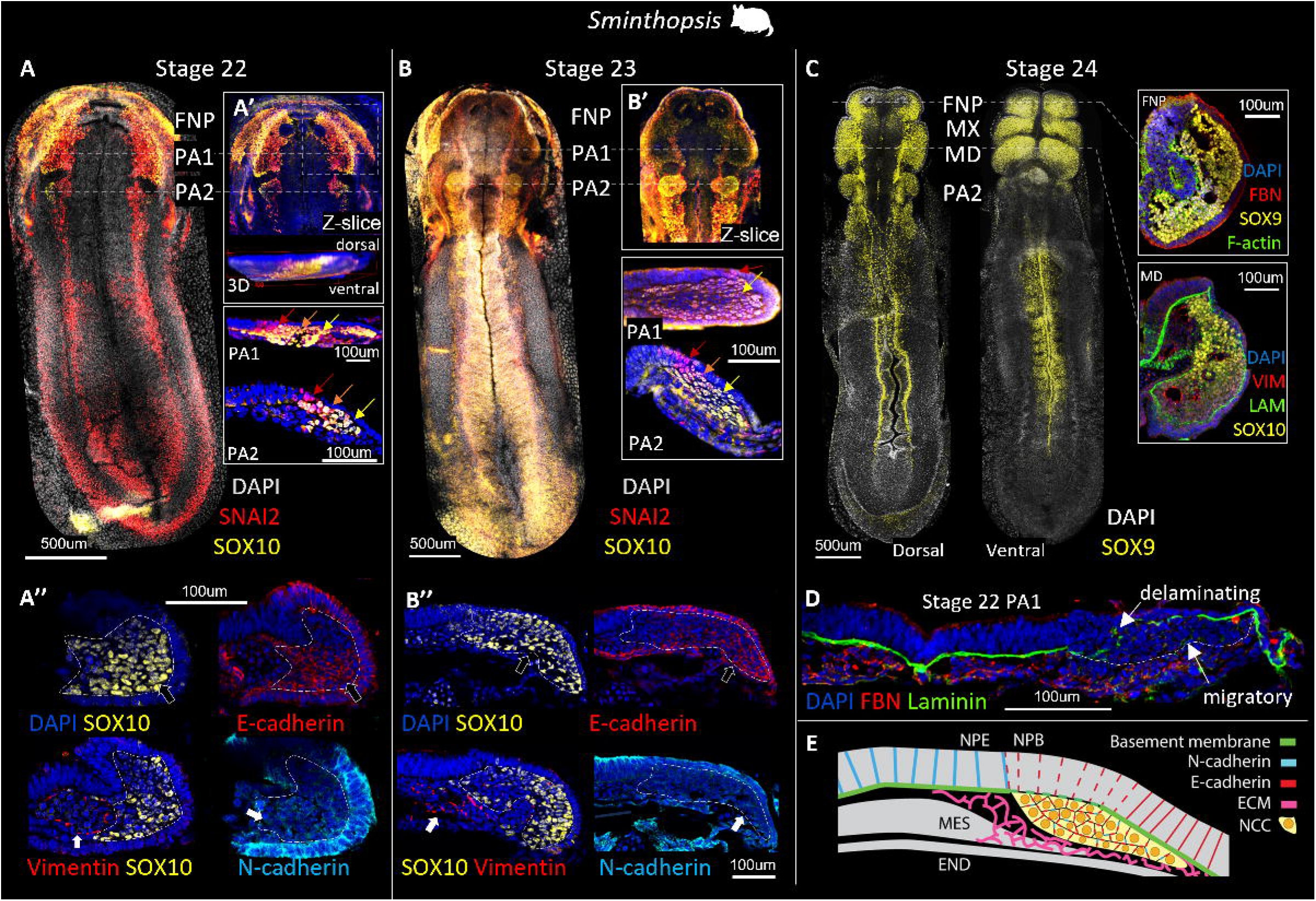
Delamination and collective migration of cranial neural crest cells. Initiation of *Sminthopsis* cranial neural crest migration across McCrady stages 22–24. **A**,**B)** Wholemount SNAI2 and SOX10 at stages 22 and 23 show continuous sheets of SOX10-positive migratory neural crest cells leaving the lateral edges of the neural plate borders, with SNAI2 repressed in delaminating cells and absent from the leader cells at the migratory front (coloured arrows); accompanying z-slices, 3D renders and PA1/PA2 tissue sections **(A⍰**,**B⍰)** show the cells displaced ventrally and expanding as densely packed, coherent sheets. **A⍰⍰**,**B⍰⍰)** Tissue sections stained for SOX10, E-cadherin, vimentin and N-cadherin: the densely packed SOX10-positive migratory cells retain strong E-cadherin junctions but lack N-cadherin and detectable vimentin, indicating retention of an epithelial-like morphology. **C)** By stage 24, SOX9-positive neural crest cells fill and demarcate the frontonasal process (FNP) and the maxillary (MX) and mandibular (MD) primordia of PA1; FNP and MD tissue sections (insets) confirm retained SOX9 with an epithelial-like junction and matrix profile. **D)** Stage 22 PA1 section showing the laminin-rich basement membrane interrupted where neural crest cells emigrate from the neural plate border, and the advancing migratory front tracking along the basement membrane of the overlying surface ectoderm and above the fibronectin-rich matrix; delaminating and migratory fronts indicated. **E)** Summary schematic of delamination and sheet migration, with the neural crest sheet wedged between the surface ectoderm and underlying mesoderm and the basement membrane interrupted at the site of emigration. NPE, neural plate epithelium; NPB, neural plate border; MES, mesoderm; END, endoderm.

To characterise the three-dimensional architecture of the migratory sheets, we performed volumetric imaging of E-cadherin and SOX10 across a series of stage 21, 22 and 23 *Sminthopsis* embryos *(48)* collected from a single litter, capturing the progression from pre-migratory to late migratory stages in temporal sequence (Figure 4). Z-stack projections, coronal z-slices, and 3D volumetric reconstructions (Figure 4D–F) together revealed that SOX10-positive leader cells migrated from the neural plate borders as a continuous, connected sheet, with retained E-cadherin-positive junctions between leading cells throughout successive stages of migration (Figure 4A’–C’). Rather than dispersing, the sheet expanded and fanned outward to sculpt the developing orofacial prominences (Figure 4A-C), with the FNP and PA1 maxillary and mandibular domains taking shape as direct extensions of the migratory sheet (Figure 3C). E-cadherin junctions were retained at the leading edge, the opposite of the progressive mesenchymalisation expected as cells move away from the neural plate (Figure 4A’’–C’’).

**Figure 4.**
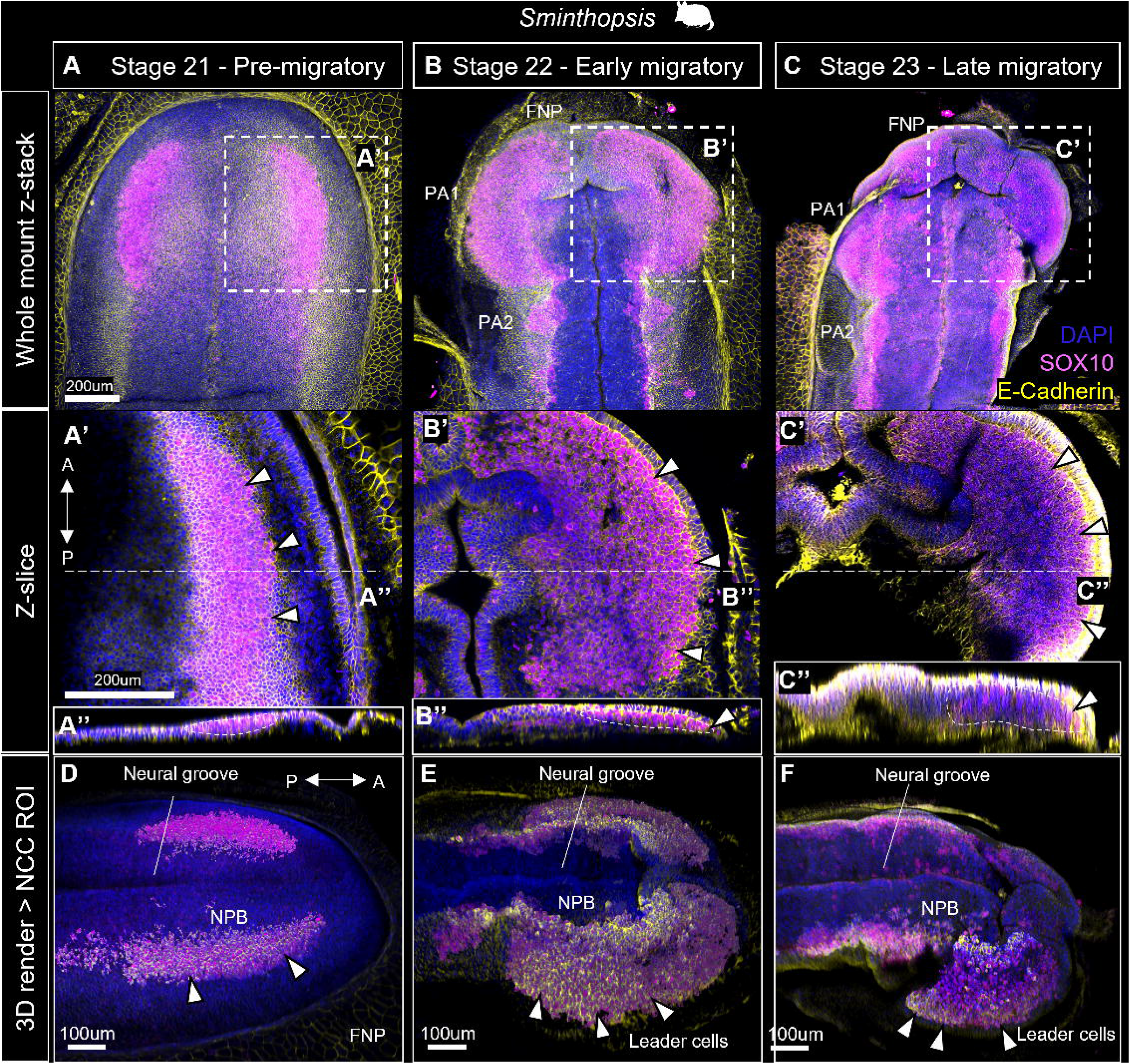
Migration of marsupial neural crest cells as an epithelial-like sheet. Wholemount SOX10 and E-cadherin imaging of *Sminthopsis* embryos from a single litter at pre-migratory (**A**, stage 21), early migratory (**B**, stage 22) and late migratory (**C**, stage 23) stages. Top panels show z-projections of the wholemount confocal stack; **A⍰–C⍰** show magnified coronal z-slices and **A⍰–C⍰** transverse z-slices, demonstrating that E-cadherin-positive junctions are retained at the leading edge of the migratory front throughout migration (arrowheads). **D–F)** 3D volumetric reconstructions of the SOX10-positive neural crest region of interest within an opaque head volume, showing the cells migrating collectively from the neural plate borders (NPB) as a connected, E-cadherin-positive sheet with leader cells at the front. A, anterior; P, posterior; NPB, neural plate border; FNP, frontonasal process.

Given the rapid formation of the orofacial prominences, we additionally asked whether increased proliferation might be the main contributor. To assess this, we quantified the proportion of mitotically active phospho-histone H3 (PH3)-positive cells within the SOX10-positive neural crest domain relative to SOX10-negative cells across the embryonic head at stages 21–23 (Figure S2). Proliferation rates were closely comparable between neural crest and non-neural crest populations (6.14% vs 6.07%; 245/3991 vs 1260/20756 cells in a single stage 22 embryo), indicating that neural crest cells are not more mitotically active than the surrounding cranial tissues at this stage. We note that this comparison is internal to the marsupial embryo and does not exclude a contribution of proliferation to the accelerated tempo relative to eutherians, which would require a cross-species comparison.

### Marsupial neural crest cells retain epithelial identity through delamination

To ask whether the cellular phenotype observed in *Sminthopsis* (and by extension *Monodelphis*) is reflected transcriptionally, we re-analysed the published *Monodelphis* single-cell embryo atlas *(47)*. The cranial neural crest trajectory was isolated from the dataset, selecting transitioning cells from the epiblast through neural plate border induction, neural crest specification, delamination, migration and ectomesenchyme based on key marker genes (Figure 5A–C). Pseudotime was fit, and cells were ordered along the trajectory (Figure 5D). Ordering of adhesion and EMT programmes across this trajectory supported a departure from the canonical amniote sequence (Figure 5E, Figure S3).

**Figure 5.**
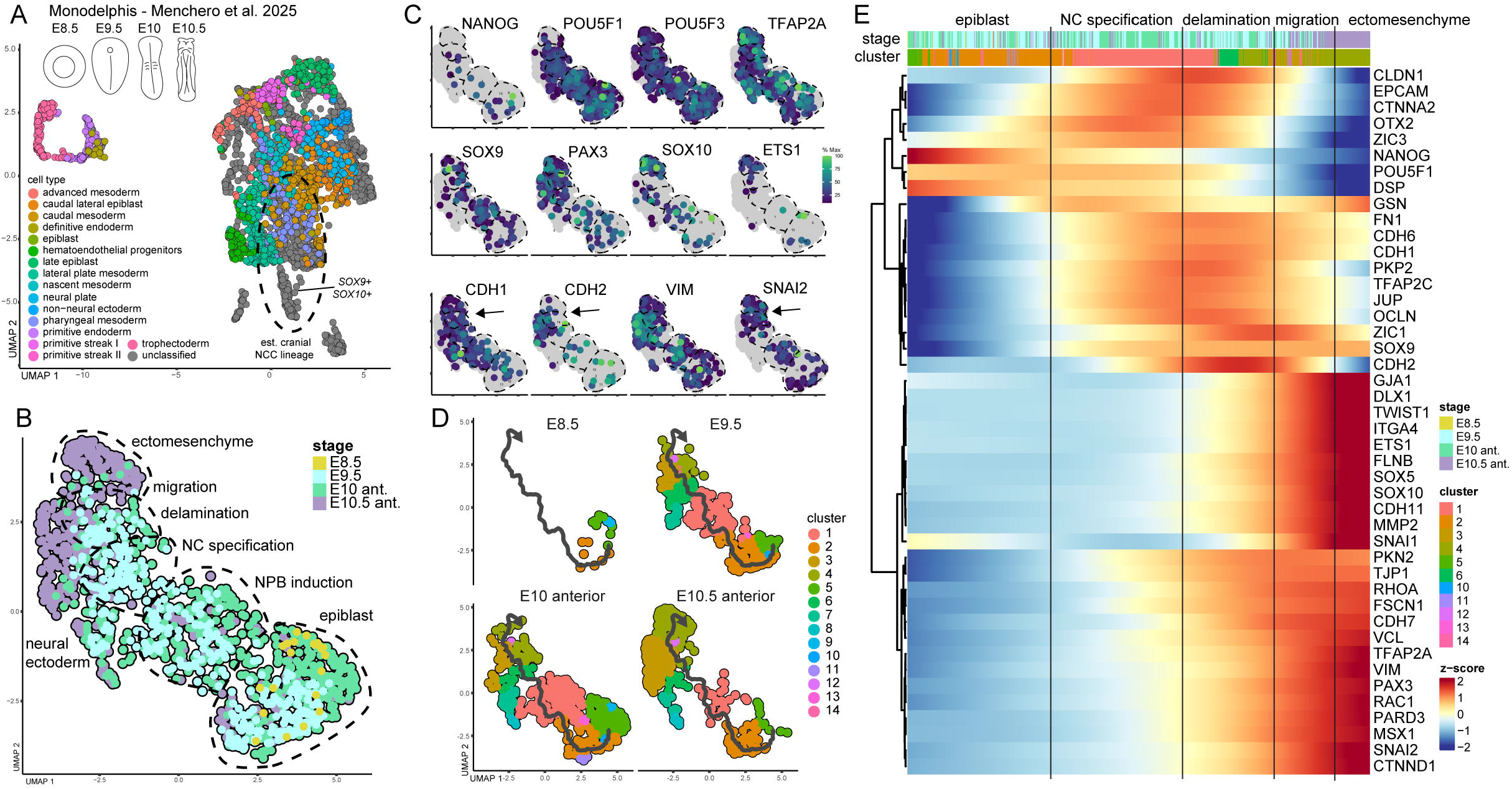
Deferred deployment of the mesenchymal programme in the marsupial cranial neural crest lineage. **A–E)** Reanalysis of the cranial neural crest lineage from the *Monodelphis* single-cell organogenesis atlas *(47)*. **A)** UMAP of the full dataset with cell-type annotations; the approximate cranial neural crest lineage (*SOX9*+/*SOX10*+) is outlined. **B)** UMAP of the isolated neural crest trajectory annotated by stage, spanning epiblast, neural plate border induction, neural crest specification, delamination, migration and ectomesenchyme. **C)** Feature plots of pluripotency and specification markers (*NANOG, POU5F1, POU5F3, TFAP2A; SOX9, PAX3, SOX10, ETS1*) and EMT-state markers (*CDH1, CDH2, VIM, SNAI2*; arrows). **D)** Trajectory coloured by stage (E8.5, E9.5, E10 anterior, E10.5 anterior). **E)** Heatmap of adhesion, cytoskeletal and EMT-associated genes across the trajectory. Epithelial junctional genes (*CDH1, EPCAM, CLDN1, OCLN, JUP, PKP2*) are induced at neural crest specification and remain high through the delamination cluster, whereas mesenchymal drivers (*TWIST1, SNAI1, MMP2, CDH11*) are induced only in the migration and ectomesenchyme clusters. CDH2 peaks at delamination. Cluster assignments as in (B).

Epithelial cadherins, namely *CDH1* (E-cadherin), *CDH6* and *CDH11* were induced during specification stages and remained elevated through delamination, migration and ectomesenchyme stages. Interestingly, in chicken, *CDH6* is active in pre-migratory neural crest and regulates EMT by driving F-actin-mediated apical cell detachment, a process tightly controlled by *SNAI2* transcriptional suppression to allow neural crest delamination and migration *(19)*. Similarly, epithelial junction genes *EPCAM, CLDN1, CTNNA1, OCLN, JUP* and *PKP2*, were induced at specification and remained high through the delamination and early/late migration clusters, whilst mesenchymal drivers *TWIST1, SNAI1, MMP2* and *CDH11* were similarly specified during delamination but rose in the migration and ectomesenchyme clusters. These contrast with mechanisms in chick and mouse, where the EMT effector programme is instead deployed before and during delamination (*15–18, 21, 51, 52*). In parallel, *CDH2* peaked at delamination and fell as cells entered migration, consistent with neuroepithelial identity and with the capacity of N-cadherin to restrain rather than drive emigration *(53)*; and *CDH11*, required for directional collective migration and contact inhibition of locomotion in *Xenopus (54)*, was induced alongside the actin regulators *FSCN1* and *FLNB (55)*, consistent with a shift toward polarised, coordinated movement. However, while these patterns largely support our cellular observations, *SNAI2, VIM* and *CDH2* transcripts showed some differences. For example, the *Monodelphis* single cell data show these genes are induced across migration and ectomesenchyme, whereas SNAI2 and vimentin protein are undetectable and E-cadherin junctions retained in migratory *Sminthopsis* cells. Because SNAI2 protein is subject to rapid post-translational turnover *(56)* and junctional E-cadherin is comparatively long-lived, we interpret the transcriptional data as indicating when the EMT programme is deployed, and the protein data as indicating whether it is functionally utilised.

Together these data suggest that marsupial cranial neural crest cells may utilize a collective migration strategy which differs from the canonical amniote sequence, with heterochronic formation of aggregates potentially acting as a reservoir of pre-migratory neural crest cells. Migratory cells at the leading edge of the aggregates were seen to repress SNAI2 after delamination, retain E-cadherin and epithelial markers, tight cell-cell contacts, lack mesenchymal markers, and migrate consistent with a collective sheet rather than as individual, dispersed mesenchymal cells. While transient E-cadherin retention has been reported in the earliest delaminating cranial neural crest cells in avian and mouse embryos before rapid mesenchymalisation *(18, 52, 57, 58)*, marsupials maintain E-cadherin and tight-contacts throughout stages of delamination and migration. This, accompanied by undetectable vimentin and SNAI2 repression, and transcriptomic support for persistent cell-cell and gap junction markers, is indicative of a partial EMT state rather than a true mesenchymal phenotype. Our findings point to a model where heterochronic marsupial neural crest cell competence is primed as early as primitive streak stages, allowing formation of pre-migratory reservoirs which in turn aid distribution of large quantities of collectively migrating cells to the rapidly forming facial prominences.

### Facial prominence patterning is most advanced in the most altricial species

We next asked whether the aggregate formation and collective migration are shared across marsupials, and whether their magnitude scales with the degree of altriciality at birth. To do this, we compared early neural crest migration across the three marsupial species differing along the altricial–precocial spectrum – the ultra-altricial (G1) *Sminthopsis*, intermediate (G2) *Monodelphis*, and the most precocial (G3) *Notamacropus* – alongside roughly stage-matched mouse (i.e. via somite number and morphological landmarks, including neural fold elevation and neural tube closure) as a eutherian comparator. Wholemount fluorescent imaging of *SOX9, SOX10, SNAI2* and *TFAP2A* in early-somite stage embryos confirmed that in each marsupial species, large accumulations of neural crest cells had already populated the presumptive FNP, PA1 and PA2 despite the neural tube remaining open (Figure 6A–C). We also observed variation in the relative extent of neural crest domain formation across the G1–G3 species. In stage 24 embryos, the jaw-forming maxillary and mandibular domains occupied a visibly greater proportion of the head in the ultra-altricial G1 Sminthopsis than in G2 *Monodelphis* or G3 Notamacropus, despite the Notamacropus embryos being considerably larger overall, whereas in roughly stage-matched mouse embryos only nascent PA1 migration and no PA2 outgrowth had occurred (Figure 6D). Because the neural crest marker available for each species differed, and because a single embryo was examined per species, we treat this as an ordinal morphological observation rather than a quantitative comparison of domain size; though corresponding area measurements are provided in Table S1, in which the frontonasal domain shows the opposite trend to the jaw-forming domains. Comparisons at the later stage 27 further highlighted the rapid outgrowth of the orofacial prominences in marsupials relative to stage-matched mouse and chicken embryos (Figure 6E–G). In the species examined, the extent of neural crest patterning of the facial prominences was greatest in the most altricial species, consistent with the hypothesis that early feeding requirements place acute developmental demands on craniofacial morphogenesis.

**Figure 6.**
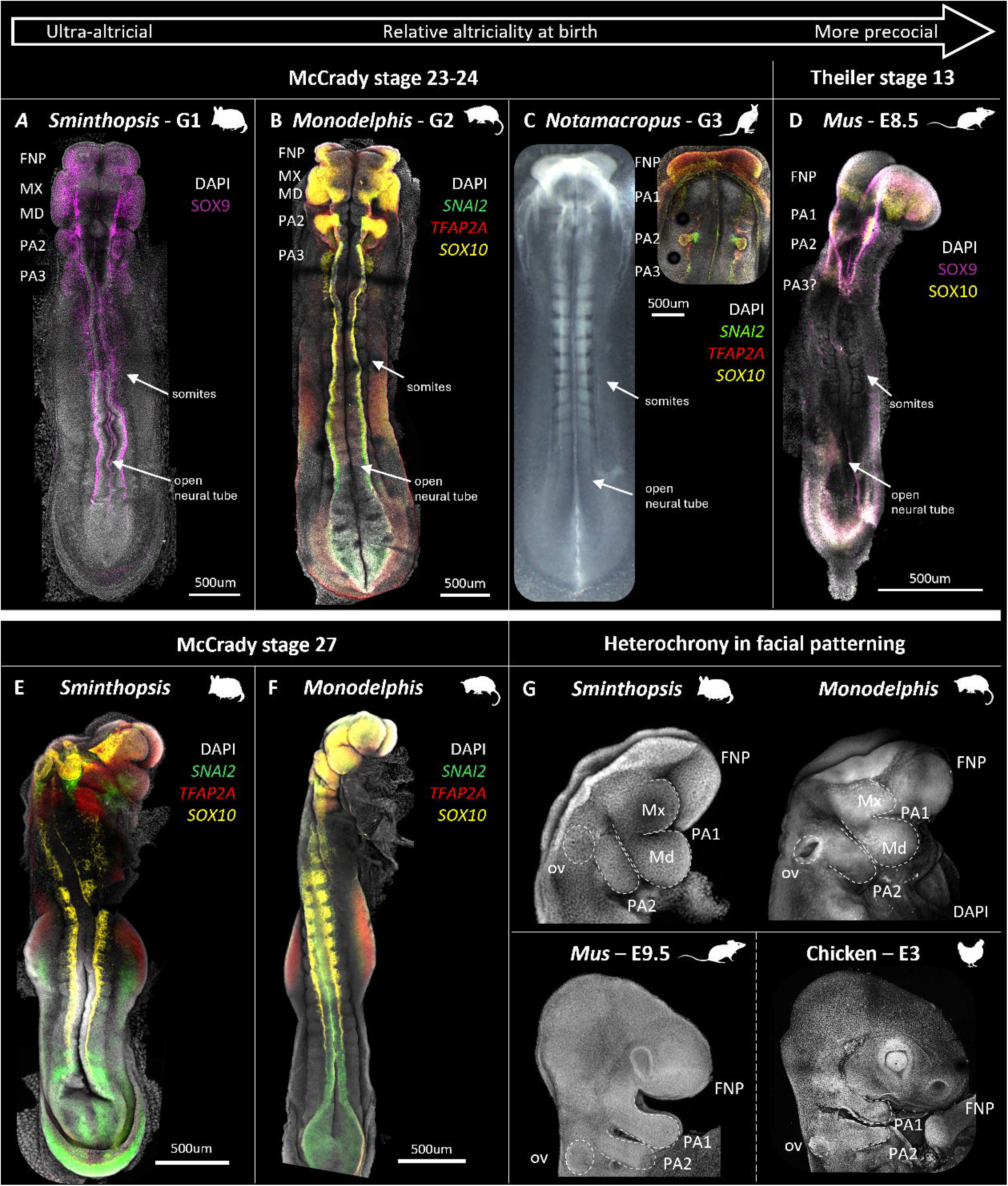
Comparative craniofacial neural crest formation across marsupials and amniotes. Neural crest formation at McCrady stage 23–24 in **A)** *Sminthopsis* (G1), **B)** *Monodelphis* (G2; *SNAI2, TFAP2A, SOX10*) and **C)** *Notamacropus* (G3), and roughly stage-matched **D)** Mus musculus (Theiler stage 13, E8.5); brightfield insets show somite number and the open neural tube. In each marsupial, large neural crest accumulations have already populated the FNP, PA1/MX–MD and PA2 despite the neural tube remaining open, whereas the stage-matched mouse shows only nascent PA1 migration and no PA2 outgrowth. **E**,**F)** Stage 27 Sminthopsis and Monodelphis showing outgrowing facial prominences. **G)** Stage 27 Sminthopsis and Monodelphis head compared with stage-matched *Mus* (E9.5) and chicken (HH18, ∼E3), emphasising the rapid outgrowth of the maxillary and mandibular primordia in marsupials. FNP, frontonasal process; MX/Mx, maxillary; MD/Md, mandibular; PA, pharyngeal arch; ov, otic vesicle.

## Discussion

In this study, we provide detailed cellular and molecular analyses of cranial neural crest formation and migration across marsupials, revealing a suite of morphogenetic properties that depart from the canonical amniote programme and instead parallel strategies described in anamniote vertebrates. The developmental mechanisms underlying marsupial craniofacial heterochrony have previously only been explored in a limited way *(45, 46, 59)*, and our results show that marsupial evolution has involved substantial plasticity in neural crest developmental programs, specifically distinct gene expression profiles and cellular trajectories compared with other amniotes (*32–34, 45*). In *Sminthopsis*, the timeline from neural crest induction to patterning and separation of the facial prominences occurs over approximately one day, based on the stage 19 to stage 24 timetable reported for this species *(48)*. This is not only faster than in *Monodelphis (45, 46)* but may be among the most rapid craniofacial patterning documented in any mammal. The timing and mode of cranial neural crest migration varies substantially even within amniotes – in eutherian mammals, cells migrate during closure of the neural folds (*14–18*), whereas in birds and reptiles migration occurs after neural tube closure *(21, 60, 61)*. Marsupial cranial neural crest cells begin migrating earlier still, before neural fold apposition and neural tube closure, from a still-planar neural plate epithelium, and we show here that this premature emigration is accompanied by three cellular behaviours not previously described in mammals: priming of the neural crest GRN and specification at primitive streak stages, heterotopic formation of dense, epithelial pre-migratory aggregates, and rapid patterning of the facial primordia as epithelial-like collective sheets.

In the canonical neural crest GRN, SOX9 both specifies neural crest identity *(51, 62, 63)* and activates *SNAI1/2* and *SOX10* to regulate the epithelial-to-mesenchymal transition, delamination, and migration of neural crest cells into the surrounding extracellular matrix *(62, 64)*. While early and persistent *SOX9* expression has previously been linked to accelerated neural crest development in *Monodelphis (45, 46)*, we also observed heterochronic activation of *FOXD3* but most notably, *SOX10* within the TFAP2A-positive neural plate borders in primitive streak-stage embryos, supporting early activation and compression of the marsupial neural crest GRN. The early co-activation of *SOX9, FOXD3, TFAP2A* and *SOX10* – spanning distinct nodes of the neural crest GRN (4, 11) – at these early stages is consistent with the neural crest specification trajectory observed in the *Monodelphis* single-cell atlas, where anterior neural crest clusters emerge and progress through specification stages earlier than in equivalent eutherian embryos *(47)*. This GRN compression, including the near-simultaneous deployment of specification and delamination-competence genes that are typically activated sequentially in other amniotes, likely underlies the precocious accumulation of the dense pre-migratory aggregates to facilitate collective migration and rapid formation of the marsupial facial prominences.

Another distinct deviation from the canonical neural crest GRN was decreased expression and localization of SNAI2 during neural crest delamination and migration. In both neural crest and disease models, SNAI2 facilitates the acquisition of mesenchymal properties by repressing E-cadherin and inducing N-cadherin (*14–18, 21, 51, 52, 56, 65*). Typically, SNAI2 is maintained in migratory neural crest cells of zebrafish, turtles and birds *(21, 66)*; in mouse, by contrast, *Snai1* rather than *Snai2* is expressed in the cranial neural crest *(66)*. In marsupial cranial neural crest however, SNAI2 is repressed rather than maintained at the onset of delamination, and E-cadherin junctions are retained in newly delaminating and migrating neural crest cells, which may contribute to their distinct collective migratory behaviour. Interestingly, these patterns are also seen in amphibians, most notably Xenopus, potentially acting via a convergent mechanism. Here, SNAI2 is repressed and E-cadherin maintained in delaminating cranial neural crest cells, which likewise exhibit collective migration of dense epithelial-like sheets *(24, 30, 67)*. Importantly however, mouse *Snai1* and *Snai2* are both dispensable for neural crest formation and delamination *(66, 68)*, indicating that the requirement for SNAI2 may vary across amniotes, and/or other EMT factors may play compensatory roles. Re-analysis of the *Monodelphis* embryo single-cell atlas *(47)* suggests the transcriptional EMT programme occurs after delamination rather than before it, so that cells leave the neural plate borders while remaining transcriptionally epithelial. Importantly though, the morphogenetic configuration we observe – i.e. SNAI2 repression, E-cadherin retention, and low vimentin in tightly packed cells across migration – is consistent with the partial EMT state recognised in collective cell migration contexts. In these cases, incomplete dissolution of epithelial identity enables cohesive group movement rather than individual dispersal *(69, 70)*, though whether this state is stabilised transcriptionally or post-translationally remains to be tested.

The heterochronic formation, heterotopic accumulation, and collective sheet migration of dense epithelial-like cranial neural crest cells observed in this study is distinct from patterns reported in other well-studied amniotes. To contextualise this finding across less studied taxa, we surveyed reports of cranial neural crest development across diverse vertebrates to define the typical formation and migration patterns represented by each clade (summarised in Table S2 and references therein). Epithelial sheet-like migration has not been previously described in eutherian mammals including rabbits, primates, rodents, and pigs, nor in birds or reptiles, although the absence of section histology in several reptile studies means the pattern cannot be excluded (Table S2). In contrast, collective migration of neural crest cell aggregates is reported in several anamniotes, including amphibians (Urodeles such as newts and axolotls, and Anurans such as Xenopus (*12, 19, 22–24, 54, 67, 71–73*) and multiple lineages of ray-finned fish *(25, 74)*. Curiously, historical descriptions of monotreme embryology additionally suggest the presence of cranial neural crest cell aggregates and sheet-like migration patterns referred to as ‘trigeminal ganglionic primordia’ *(35, 36, 75)*, though this lacks molecular support.

When cranial neural crest migratory strategies are further mapped across vertebrates, these different strategies appear to align with life history rather than phylogenetic position (Figure 7). In many anamniote species, heterochronic acceleration and collective migration of epithelial-like cranial neural crest cells enable the early formation of skeletal feeding and respiratory structures in altricial larvae *(25, 74)*. The functional requirements for a well-formed orofacial apparatus at early, altricial life stages in marsupials may have promoted the evolution of similar neural crest migration strategies to those utilised in anamniotes, representing characteristics that have evolved independently in response to convergent life history constraints. The presence of ‘trigeminal ganglionic primordia’ in monotremes raises the intriguing possibility that neural crest cell aggregates and collective sheet migration may in fact represent an ancestral mammalian strategy, later lost in eutherian mammals (Figure 7). Whilst speculative, monotremes, like marsupials, produce altricial young and continue development ex utero through extended lactation *(35, 75, 76)*, potentially facing comparable early feeding constraints.

**Figure 7.**
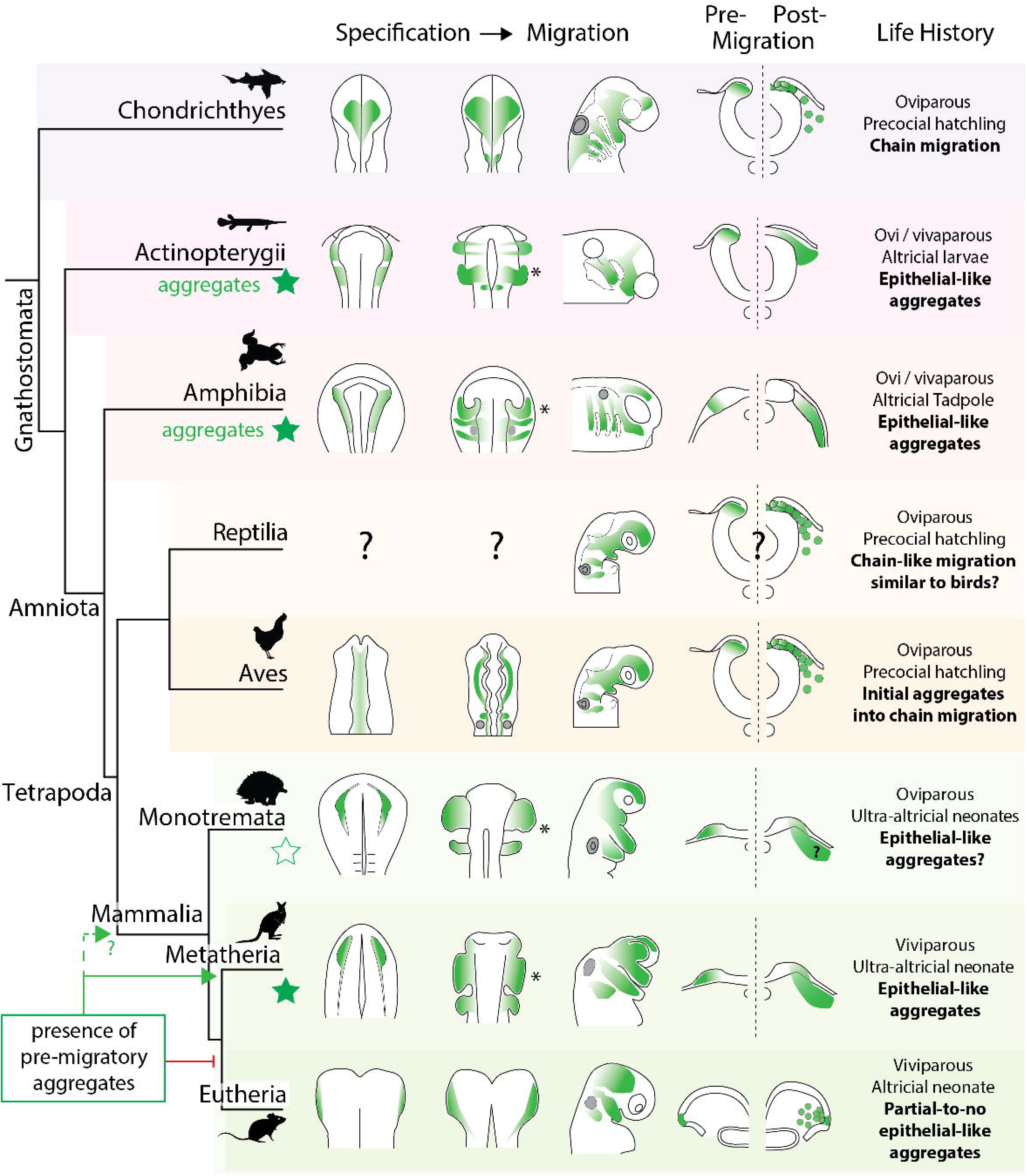
Tracing the mode of cranial neural crest migration across vertebrates. Vertebrate phylogeny showing documented patterns of cranial neural crest migration alongside life history and mode of reproduction, with neural crest cells and their migration shaded green and mapped across specification, migration, and pre- and post-migratory stages. Collective migration of epithelial-like sheets (stars) has evolved in ray-finned fishes (Actinopterygii), amphibians and marsupials (Metatheria), and is suggested in monotremes (Monotremata) — lineages producing altricial young with immediate independent feeding requirements — whereas oviparous sharks (Chondrichthyes), reptiles, birds (Aves) and studied eutherian mammals produce more precocial young and migrate distinct streams of mesenchymal neural crest cells. Question marks denote clades for which the pattern is inferred rather than directly documented. This distribution is consistent with epithelial-like neural crest aggregates representing an ancestral state that was retained or re-emerged in altricial-feeding lineages (green arrow) and was reduced or lost in eutherians (red bar) with the evolution of longer gestation and precocial young.

The evolution of pre-migratory aggregates in marsupials may be a functional adaptation to facilitate their accelerated craniofacial development *(34, 38, 43)*. Cranial neural crest aggregates may act as a reservoir to give a developmental “head start”, supplying increased numbers of progenitors to the orofacial prominences. The number of neural crest cells allocated to the orofacial prominences is known to regulate species-specific differences in craniofacial morphology *(77)*, and neural crest cells are known to control the timing of differentiation as well as the structural and functional integration of the craniofacial musculoskeletal system *(78)*. As such, these results provide a plausible cellular explanation for how marsupials achieve accelerated formation of the craniofacial complex to support the demands of early feeding (32–34). The pre-migratory aggregates function as a stockpile of migration-competent progenitors, and the collective sheet migration mode delivers this expanded progenitor pool rapidly and coherently to the facial prominences. Additional support for this hypothesis comes from comparative analyses of facial patterning across G1– G3 marsupial groups (Figure 6), where altricial marsupials possess more outgrowth of the orofacial prominences, as first described by Hill and Watson *(32)*.

Together, our observations suggest a plausible cellular mechanism by which altricial feeding structures may evolve across diverse taxa, and highlight cranial neural crest heterotopy and collective migration as potential drivers of adaptive craniofacial evolution. More broadly, these marsupial-specific patterns illustrate how evolutionary changes in developmental timing can reshape neural crest cell behaviour to meet life history demands, with implications for understanding the cellular and molecular plasticity underlying craniofacial diversity across vertebrates.

### Limitations of the study

This study is a comparative morphological description based on embryos of three marsupial species that cannot be obtained in large numbers, and several limitations follow from that. First, all observations derive from fixed material: we describe the configuration, adhesion profile and spatial progression of cranial neural crest cells across a staged series, but we do not observe cell movement directly. Our inference of collective behaviour therefore rests on the retention of cell–cell contacts and on the coherence of the migratory mass across successive stages, not on live imaging or cell tracking, and we cannot exclude that individual cells move independently within an apparently cohesive sheet. Second, timed marsupial embryos are available in small numbers and cannot be generated on demand; several panels represent a single embryo, and the comparative observations in Figure 6 derive from one stage 24 embryo per species and are presented as ordinal rather than quantitative. Third, we have not perturbed the pathway: we have no functional test of whether E-cadherin retention or SNAI2 repression is required for the migratory mode we describe, and such experiments are not currently tractable in these species. Fourth, the single-cell re-analysis orders transcriptional events relative to one another within a pseudotemporal trajectory, and does not establish absolute developmental timing nor independently establish that cells assigned to a given cluster have physically delaminated. Finally, the comparative survey in Table S2 reflects our reinterpretation of published images across taxa studied with heterogeneous methods, and the migratory mode of several clades remains inferred rather than directly documented.

## Methods

### *Sminthopsis* Husbandry and Embryo Collection

*Sminthopsis crassicaudata* (Dunnarts) were obtained from an experimental colony run within the School of Biosciences. Female and male dunnarts were allocated into mating pairs in a 3:1 or 2:1 ratio of females to males. Detection of pregnancy for timed embryo collection was performed using recently described methods *(48)*, where paired females were monitored for weight changes as the primary indicator of pregnancy, and females which showed consecutive days of weight increase after ovulation were used for embryo collection. Dunnarts were humanely killed by cervical dislocation; the paired uteri were then removed from the peritoneum and opened in DEPC-PBS. Pre-implantation embryos (<stage 28) were rolled out of the uterus, while implanted embryos were carefully cut out with their membranes intact. Embryos were transferred into 4% paraformaldehyde (PFA) and left to fix at 4°C overnight or over the weekend.

All animal experiments were performed under University of Melbourne animal ethics numbers 10206 and 26863, and conformed to the Australian National Health and Medical Research Council (2013) guidelines. Embryos were collected between McCrady stages 18 and 27, prior to gonadal differentiation, and sex was not determined; the influence of sex on the reported phenotypes could therefore not be assessed.

### *Monodelphis* and Tammar Wallaby Embryos

Fixed *Monodelphis domestica* embryos were supplied from the University of California, Los Angeles, approved by the UCLA Institutional Animal Care and Use Committee.

Fixed Tammar wallaby *Notamacropus (Macropus) eugenii* embryos were supplied from a breeding colony of the University of Melbourne and conformed to the Australian National Health and Medical Research Council (2013) guidelines.

### Mouse and chicken embryos

Mus musculus embryos were obtained from the University of Melbourne Animal Facility under approved application 26868, and conformed to the Australian National Health and Medical Research Council (2013) guidelines. Embryos were staged by somite count according to Theiler *(79)*.

Fertilised chicken eggs were obtained from Specialized Breeders Australia, incubated at 37.5 °C and staged according to Hamburger and Hamilton *(80)*.

### Whole-mount Fluorescent Imaging

Gene expression analysis was performed using immunofluorescence and hybridization chain reaction (HCR) *(81)*. In both cases, whole embryos fixed in PFA were washed in RNAse free 1X PBS with 0.1% Tween-20 (PBST) and then dehydrated in a 25%, 50%, 75%, and 100% methanol series (in PBS) on ice. Embryos were then stored in 100% methanol at -20°C until use. Target genes were chosen to represent key nodes of the neural crest cell gene regulatory network, including pluripotency (*POU5F1*), initiation (*FOXD3, TFAP2A*), specification (*SOX9, SNAI2* (*SLUG*)), migration (*SOX10*), as well as markers for general morphology (DAPI, F-actin/phalloidin) and for cell adhesion and extracellular matrix (E-cadherin, N-cadherin, vimentin, fibronectin and laminin; Table S3).

HCR probes (Molecular Instruments, Inc., Los Angeles, CA) were initially designed against target gene sequences from the *Sminthopsis crassicaudata* transcriptome *(82)*. However, to ensure cross-reactivity between marsupials, probe sequences were BLASTed and retained only if they were specific to the gene of interest and had >85% similarity to *Monodelphis* (the most distantly related species examined here). HCR was performed using the protocol provided by Molecular Instruments *(81)* with minor modifications based on the stage of the embryos. Briefly, embryos were rehydrated with the reverse methanol series (75%, 50%, 25% methanol, then 100% DEPC-PBST) on ice. Embryos were then incubated with proteinase-K at room temperature for 5 minutes (stages 24-25) or 10 minutes (for stages 27-29), then post-fixed in DEPC-PFA for 20 minutes at room temperature. Embryos at stages prior to stage 24 were not incubated with proteinase-K and were not post-fixed. All embryos were incubated with 10-30pmol of probes for *POU5F1, SNAI2, FOXD3, TFAP2A, SOX9* and/or *SOX10* in hybridisation buffer at 37°C overnight. On the second day, embryos were washed in wash buffer and DEPC-SSCT and were then incubated with 30pmol of H1 and H2 hairpins in amplification buffer at room temperature overnight. On the third day, embryos were washed in DEPC-SSCT and incubated with DAPI (1:10,000 in PBST), before being washed again in PBST.

Methanol dehydrated embryos were used for immunofluorescence but were not rehydrated in a series. Instead, embryos were transferred directly from 100% methanol into blocking buffer (1X PBS with 0.1% Tween-20 (PBST) with 3% BSA) and left to block for at least 1 hour, rotating at 4°C. Primary antibodies (Table S3) were diluted in PBST with 1% BSA and incubated with embryos overnight at 4°C while rotating. On the second day, embryos were washed with PBST to remove any unbound primary antibodies, then incubated with secondary antibodies (diluted at 1:500 in PBST with 1% BSA) at room temperature overnight in the dark while rotating. On the third day, embryos were washed in PBST and then incubated with DAPI (1:10,000 in PBST), before a final wash in PBST prior to mounting.

### Mounting of Embryos for microscopy

Mounting was performed based on the size, stage and shape of the embryo. Mounting of early-stage embryos (stage 19-22) involved positioning the embryo flat on the slides with the dorsal surface facing upwards. Embryos often adopted the curvature of the vesicle and occasionally required further flattening through cutting the extra embryonic membranes. Embryos were then overlaid with glycerol mounting media (90% glycerol in 0.1 M PBS with 50 mg/ml propyl gallate) and a coverslip gently applied. Mounting of stage 23+ embryos required generation of custom silicon wells using aquarium-grade sealant. Embryos were positioned within the well and filled with the same fluorescent mounting media. A cover slip was then placed on top to seal the embryo within the chamber.

### Tissue Section Immunofluorescence

Whole embryos were transferred to 30% sucrose in PBS and allowed to sink, typically overnight. Embryos were then positioned within cryomolds in Tissue-Tek O.C.T. mounting media (ProSciTech) and stored at −80 °C until use. 15–20 µm cryosections were cut and placed on alternating superfrost slides for successive immunostaining. Slides were washed in 1% Triton-X in 1X PBS (PBTX) and then blocked for at least 1 hour with 2% BSA in PBS at room temperature in a humidified chamber to prevent drying. Slides were then washed in PBS again and then incubated with primary antibodies diluted in PBS at 4°C overnight. On the second day, sections were washed in PBS and incubated with secondary antibodies (1:500 in PBS containing 1% BSA) at room temperature in the dark for 1 hour. Slides were then washed in PBS and incubated with DAPI (1:10,000 in PBS), before being washed again and mounted with glycerol mounting media and a coverslip.

### Microscopy and post-processing

Embryos and slides were imaged on a Nikon A1R confocal microscope with NIS-Elements software. Whole mount scans were captured with a 10x PL APO Lambda MRD00105 air objective (NA = 0.45), while tissue section images were imaged with a 40x PL FLUO MRH01401 oil objective (NA = 1.3). The number of embryos, litters and sections underlying each panel is stated in the corresponding figure legend; panels representing a single embryo are identified as such.

### Image processing and quantification

All image post-processing was performed using ImageJ (Fiji) for visualization, z-projection or z-slicing; or Imaris (Oxford Instruments, UK) for 3D volumetric reconstruction and quantification.

Confocal imaging data was imported into Imaris v10.2 (Oxford Instruments, UK) for volumetric visualization and surface rendering. Intensity thresholding of SNAI2 and SOX10 expression channels were used to create surfaces for 3D volumetric rendering. To calculate relative channel intensities between cells, the Imaris “cells” module was used to segment cells via DAPI signal (low threshold to capture cells broadly) and generate cell counts/data across the embryonic head encompassing the headfolds. Cells data was exported from Imaris in CSV format, imported into R studio for further analysis and graphing. The average channel intensities per cell for each channel were plotted against cell position in the frontal axis of the embryo.

Cell morphology and proliferation were determined within neural crest cells in Imaris via creation of a region of interest and channel mask. Initially, neural crest cells were identified using SOX10-positive signal in the Imaris “spots” module to generate a surface of the migratory neural crest cell domain. To visualize the migratory mass without interference from the overlying ectoderm or surrounding tissues, a mask was created for each channel from the surface, before the background volume made opaque. This allowed specific visualization of cell junctions between neighbouring neural crest cells, with E-cadherin signal from the overlying ectoderm. To determine proliferation rates between neural crest cells and non-neural crest cells, the neural crest surface was used to generate a neural crest-specific and non-neural crest masks. The ‘cells’ function was then used on the DAPI channel of each mask to calculate the number of cells within the region of interest, followed by the PH3 channel to calculate the number of proliferating cells. We then calculated the proportion of proliferative cells (proliferative/total*100) within a region to determine the gross proliferation rate.

To measure the relative area of FNP & PA1-3 neural crest cell domains between marsupials, area of each domain was calculated from the z-projection images of stage 24 embryos using ImageJ (Fiji). Domain boundaries were defined using the neural crest marker available for each species (SOX9 in S. crassicaudata, *SOX10* in *M. domestica* and *N. eugenii*). Because these markers label partially distinct subsets of the neural crest population, and because one embryo was measured per species, the resulting proportions are reported in Table S1 as descriptive values and are not used to support a quantitative comparison of domain size. The total head area was defined as the anterior-most edge of the FNP domain to immediately below PA2, and each neural crest cell-domain was calculated using neural crest cell-gene expression to define the domain boundaries. Total shape was captured using the Polygon-selection tool and area calculated using the Measure function. Neural crest domain area was divided by total head area to obtain a % proportion of total head area (Table S1).

### Single-cell RNA-sequencing re-analysis

The published *Monodelphis* domestica embryo single-cell atlas *(47)* was obtained from the Gene Expression Omnibus (GSE264385), comprising processed cells across embryonic day E8.5, E9.5, E10.0 (anterior) and E10.5 (anterior). Analysis was performed in Bioconductor packages and Monocle3 *(83)* in Rstudio. Cells were clustered without supervision (Leiden; Monocle3 cluster_cells), and clusters were assigned to the cranial neural crest lineage on the basis of marker gene expression visualised on the embedding (SOX9, SOX10, FOXD3, TFAP2A); no marker gene was used to order cells along the trajectory. Pseudotime was fitted using Monocle3 with the root anchored in POU5F1 expressing E8.5 cells. For the heatmap in Figure 5E, expression was normalized across pseudotime and scaled to a z-score per gene. Because pseudotime is internally normalised, it orders transcriptional events relative to one another but does not provide an absolute developmental timescale, and no cross-species timing comparison is made from these data.

## Acknowledgements

The authors would like to thank The School of BioSciences (University of Melbourne) animal facility staff for the daily management of the Dunnart colony, particularly Shiralee Whitehead. We thank staff from the University of Melbourne Biological Imaging platform (BOMP) for microscopy usage and imaging assistance.

## Author contributions

AHN and AJP conceived the study. AHN and ERF performed the experiments. AHN reconstructed and analysed imaging data. AHN and ATM performed image quantification and plotting. AHN, ERF, JCH and BML monitored dunnarts for pregnancy and collected embryos. AMC and KES supplied opossum embryos. GS and MBR supplied tammar wallaby embryos. SO contributed to project administration and resources. AHN, ERF, RAS, YW and AJP wrote the manuscript. All authors reviewed the data and gave final approval of the manuscript.

## Data availability

The single-cell RNA-sequencing dataset re-analysed in this study is publicly available from the Gene Expression Omnibus under accession GSE264385 *(47)*. Confocal imaging data supporting the findings of this study are presented in the manuscript figures.

## Figure Legends

**Figure S1.Monodelphis collective migration.**

Whole mount and optical sections of *SNAI2* and *SOX10* expression, highlighting collective-like migratory properties of cranial neural crest cells

**Figure S2. Proliferation within the neural crest domain is comparable to that in surrounding cranial tissue.**

Extension of Figures 3 and 4. Pre-migratory (stage 21), early migratory (stage 22) and late migratory (stage 23) *Sminthopsis* embryos stained for SOX10 and phospho-histone H3 (PH3). Top panels show z-projections of the wholemount confocal stack; middle panels show coronal z-slices; bottom panels show 3D volumetric reconstructions of the SOX10-positive neural crest region of interest within an opaque head volume. Quantification of the early-migratory (stage 22) domain shows no increase in the proportion of PH3-positive (mitotically active) cells within the SOX10-positive neural crest population relative to SOX10-negative cells across the head (6.13% vs 6.07%). A, anterior; P, posterior; NPB, neural plate border; ROI, region of interest.

**Figure S3. Extended single-cell expression of the *Monodelphis* neural crest trajectory.**

Extension of Figure 5. Additional UMAP feature plots for adhesion, cytoskeletal, EMT-state and specification genes across the *Monodelphis* cranial neural crest trajectory not shown in Figure 5C, supporting the deferred deployment of the mesenchymal programme described in the main text.

