## Supplementary material for "Craniofacial heterochrony in marsupials involves dynamic changes to cellular and molecular mechanisms underlying neural crest migration": Figure S1-S3

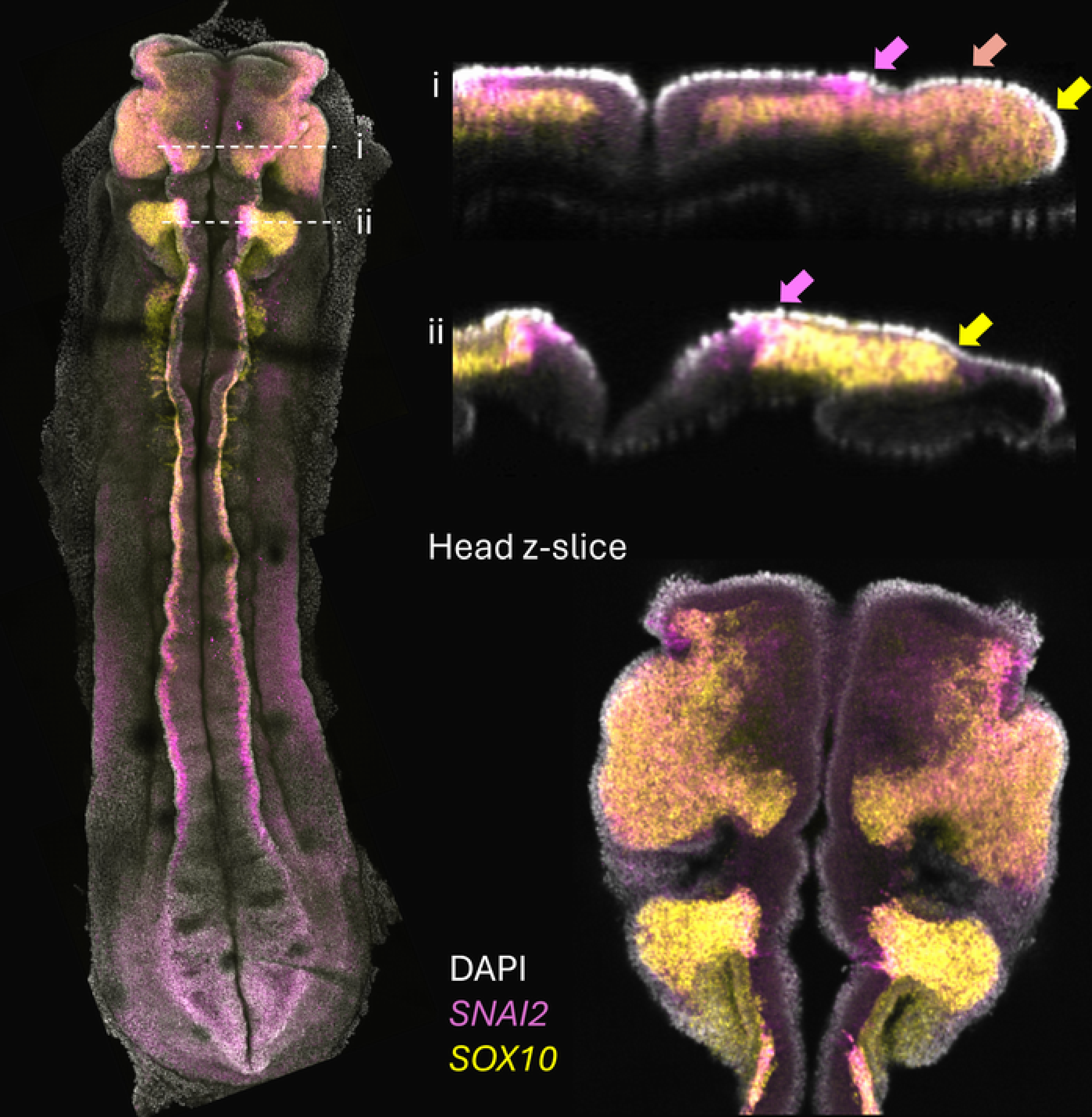

i
ii
i
ii
Head z-slice
DAPI
*SNAI2*
*SOX10*

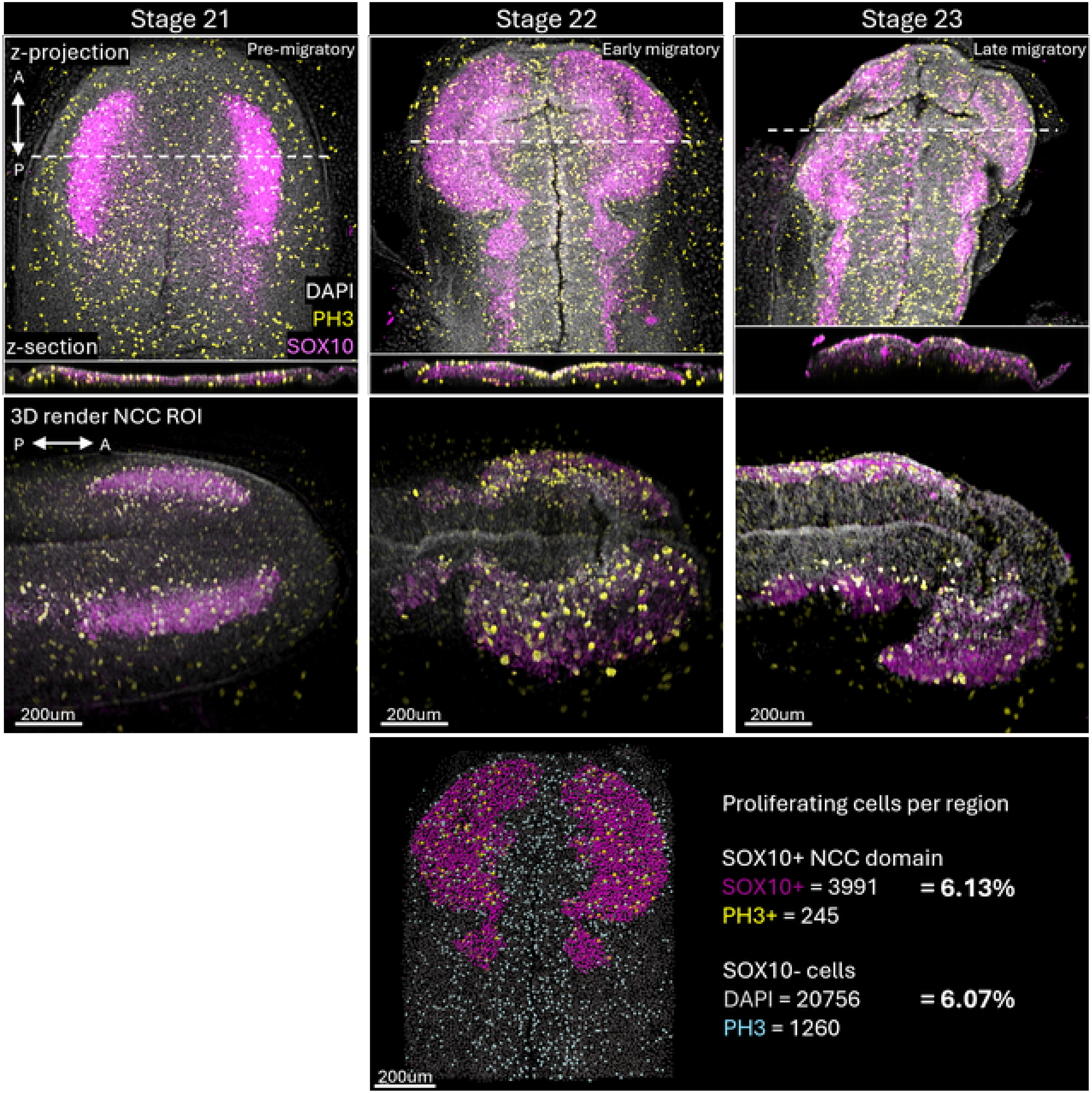

Stage 21
Stage 22
Stage 23
z-projection
Pre-migratory
Early migratory
Late migratory
A
P
DAPI
PH3
SOX10
z-section
3D render NCC ROI
P
A
200um
200um
200um
200um
Proliferating cells per region
SOX10+ NCC domain
SOX10+ = 3991
= 6.13%
PH3+ = 245
SOX10- cells
DAPI = 20756
= 6.07%
PH3 = 1260

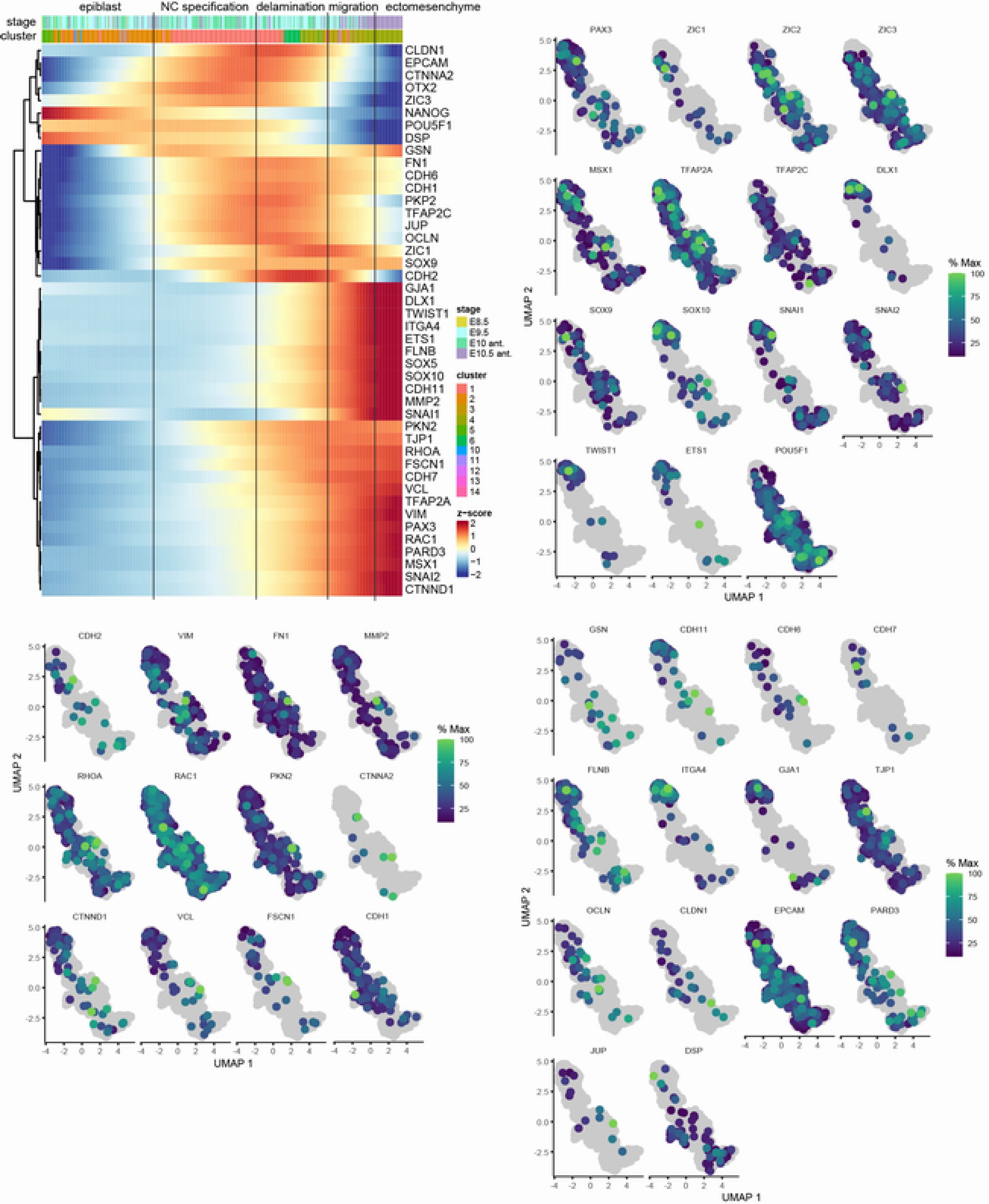

epiblast
NC specification
delamination
migration
ectomesenchyme
stage
cluster
CLDN1
EPCAM
CTNNA2
OTX2
ZIC3
NANOG
POU5F1
DSP
GSN
FN1
CDH6
CDH1
PKP2
TFAP2C
JUP
OCLN
ZIC1
SOX9
CDH2
GJA1
DLX1
TWIST1
ITGA4
ETS1
FLNB
SOX5
SOX10
CDH11
MMP2
SNAI1
PKN2
TJP1
RHOA
FSCN1
CDH7
VCL
TFAP2A
VIM
PAX3
RAC1
PARD3
MSX1
SNAI2
CTNND1
stage
E8.5
E9.5
E10 ant.
E10.5 ant.
cluster
1
2
3
4
5
6
10
11
12
13
14
z-score
2
1
0
−1
−2
PAX3
ZIC1
ZIC2
ZIC3
MSX1
TFAP2A
TFAP2C
DLX1
SOX9
SOX10
SNAI1
SNAI2
TWIST1
ETS1
POU5F1
% Max
100
75
50
25
UMAP 2
UMAP 1
CDH2
VIM
FN1
MMP2
RHOA
RAC1
PKN2
CTNNA2
CTNND1
VCL
FSCN1
CDH1
GSN
CDH11
CDH6
CDH7
FLNB
ITGA4
GJA1
TJP1
OCLN
CLDN1
EPCAM
PARD3
JUP
DSP
